## Supplemental Information for "Integrated omics networks reveal the temporal signaling events of brassinosteroid response in *Arabidopsis*"

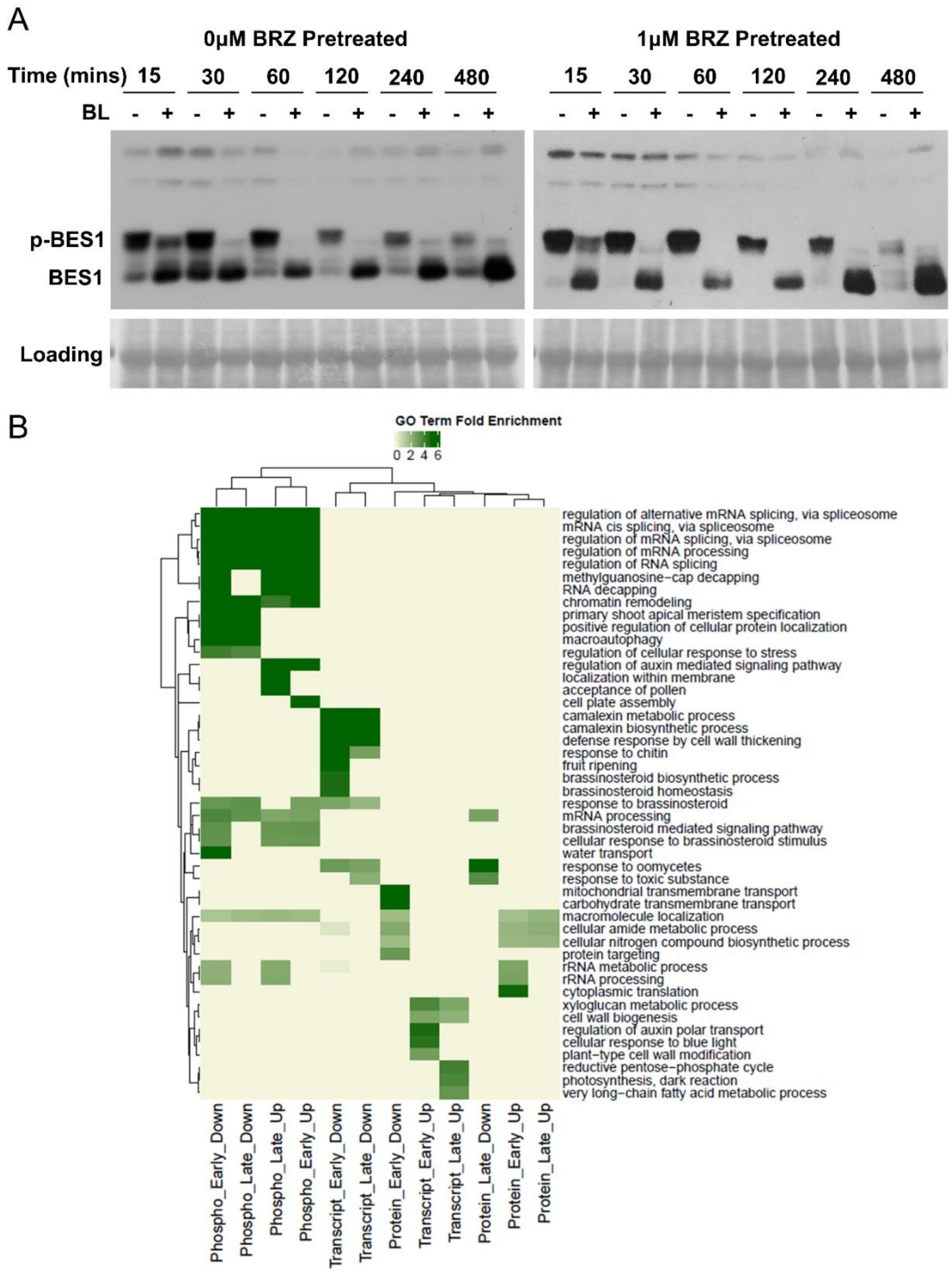

**Fig S1.** Effect of BL treatment on transcript, non-modified protein, and phosphorylated protein levels. (A) Western blot of phosphorylated (p-BES1) and non-modified BES1 protein after BL treatment. (left) No BRZ pre-treatment. (right) BRZ pre-treatment. (B) Heatmap of GO terms enriched for different omics data (transcript, protein, phospho) at different times (early, 1 hour or earlier after BL treatment) (late, after 2 hours of BL treatment).

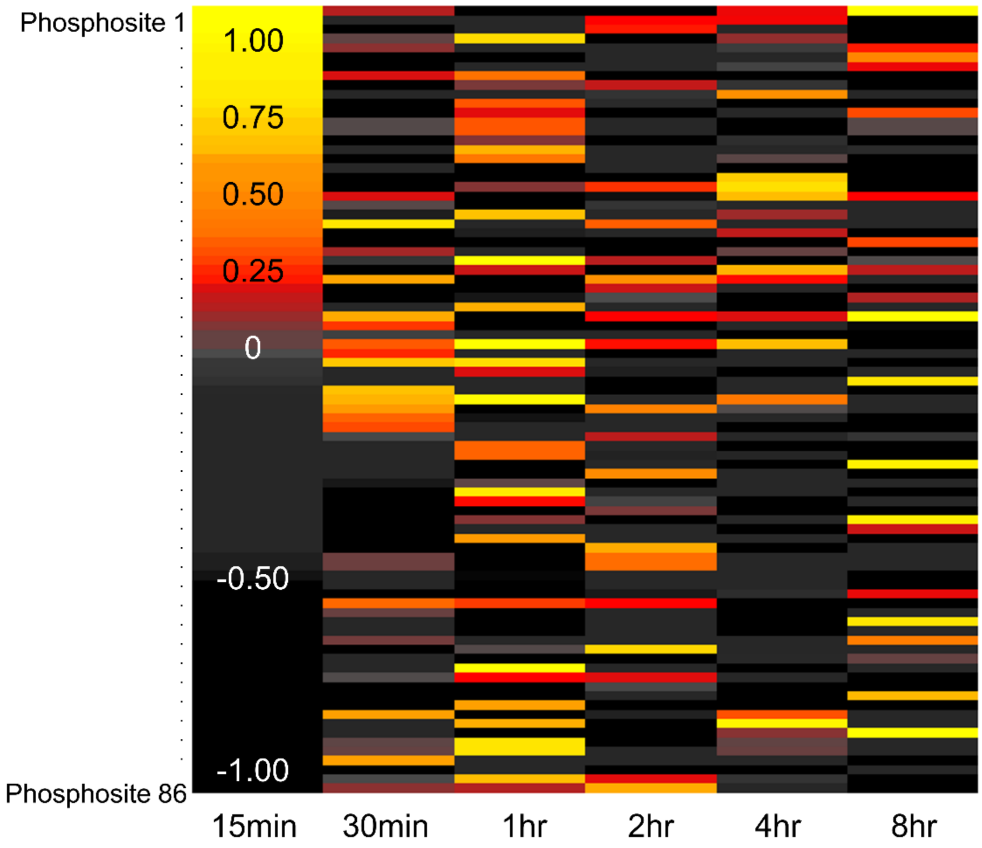

**Fig S2.** Correlation between non-modified protein abundance and phosphosite abundance for kinases DE in response to BR. We identified 86 phosphosites as p-loop activation domains that were DE in response to BR. We then correlated the protein abundance with the phosphosite intensity values for these 86 sites, which are shown in this heatmap. Yellow represents the highest positive correlation, orange is moderate correlation, and red to black is low to no correlation. Phosphosites are rows, time points are columns.

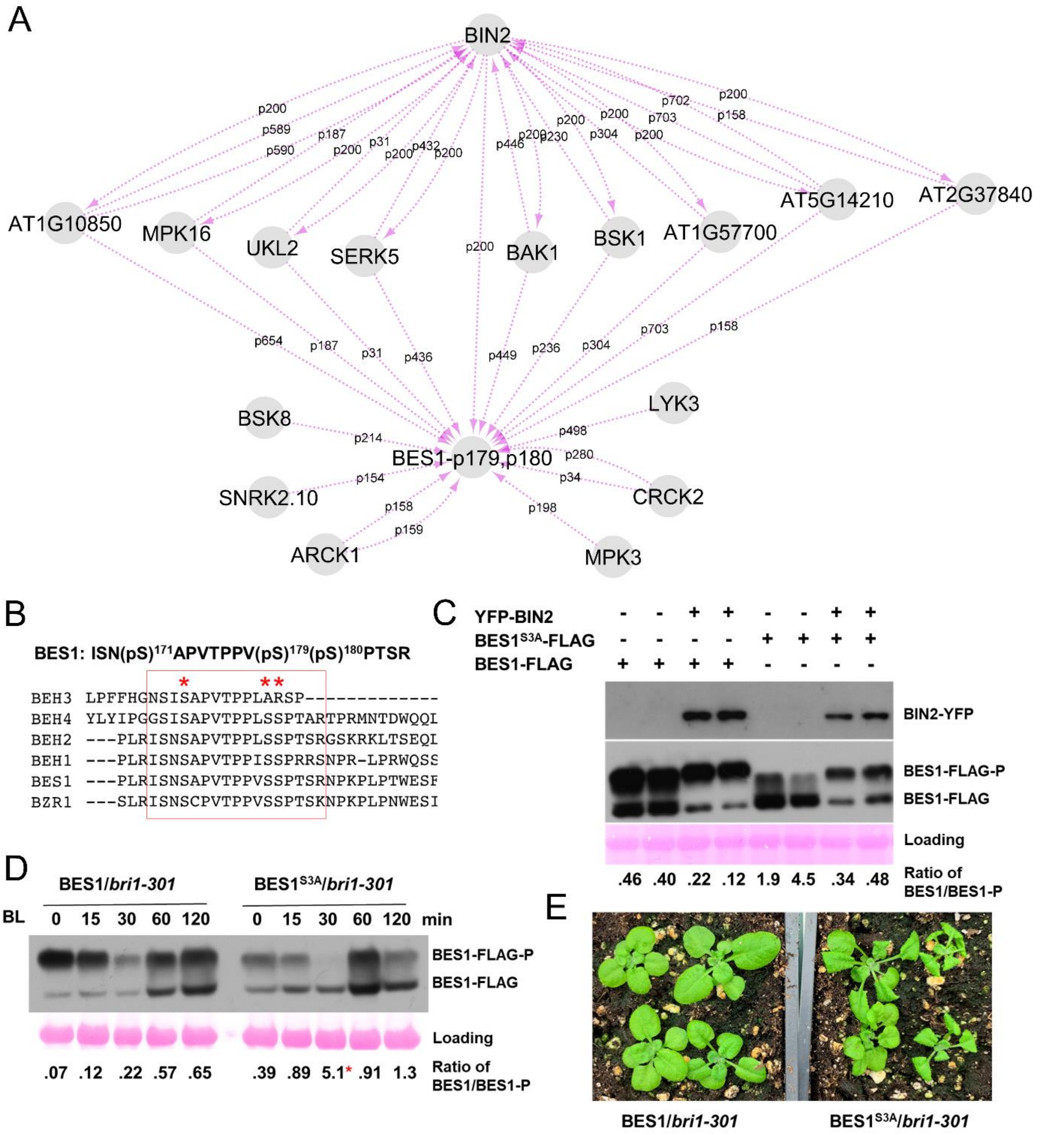

**Fig S3.** Network prediction and experimental validation that S171, S179, and S180 on BES1 are BIN2-phosphorylation sites (A) Kinase-signaling network prediction for the doubly-phosphorylated form of BES1, which is phosphorylated on sites 179 and 180. Edge labels represent the phosphosite used to predict the regulation. (B) Conservation of S171, S179, and S180 in BES1 with its homologs. Protein alignment was performed using Clustal Omega (https://www.ebi.ac.uk/Tools/msa/clustalo/) . (C) Transient expression of *BES1-FLAG* and *BES1$^{S3A}$-FLAG* in *Nicotiana benthamiana* leaves, with or without BIN2-YFP (n=2 biological replicates for each combination). (D) Phosphorylation status of BES1-FLAG and BES1$^{S3A}$-FLAG in response to BL treatment. n=12 T1 seedlings were used for each treatment. The * indicates that the ratio may not be accurate due to very low level of phosphorylated BES1 in this sample. (E) Growth phenotype of 3-week-old T1 transgenic plants of BES1-FLAG (left) and BES1$^{S3A}$-FLAG (right) in *bri-301* mutant background. n=18 out of 24 BES1$^{S3A}$-FLAG plants showed the BR-gain-of-function growth phenotype. Each T1 plant represents an independent transgenic line.

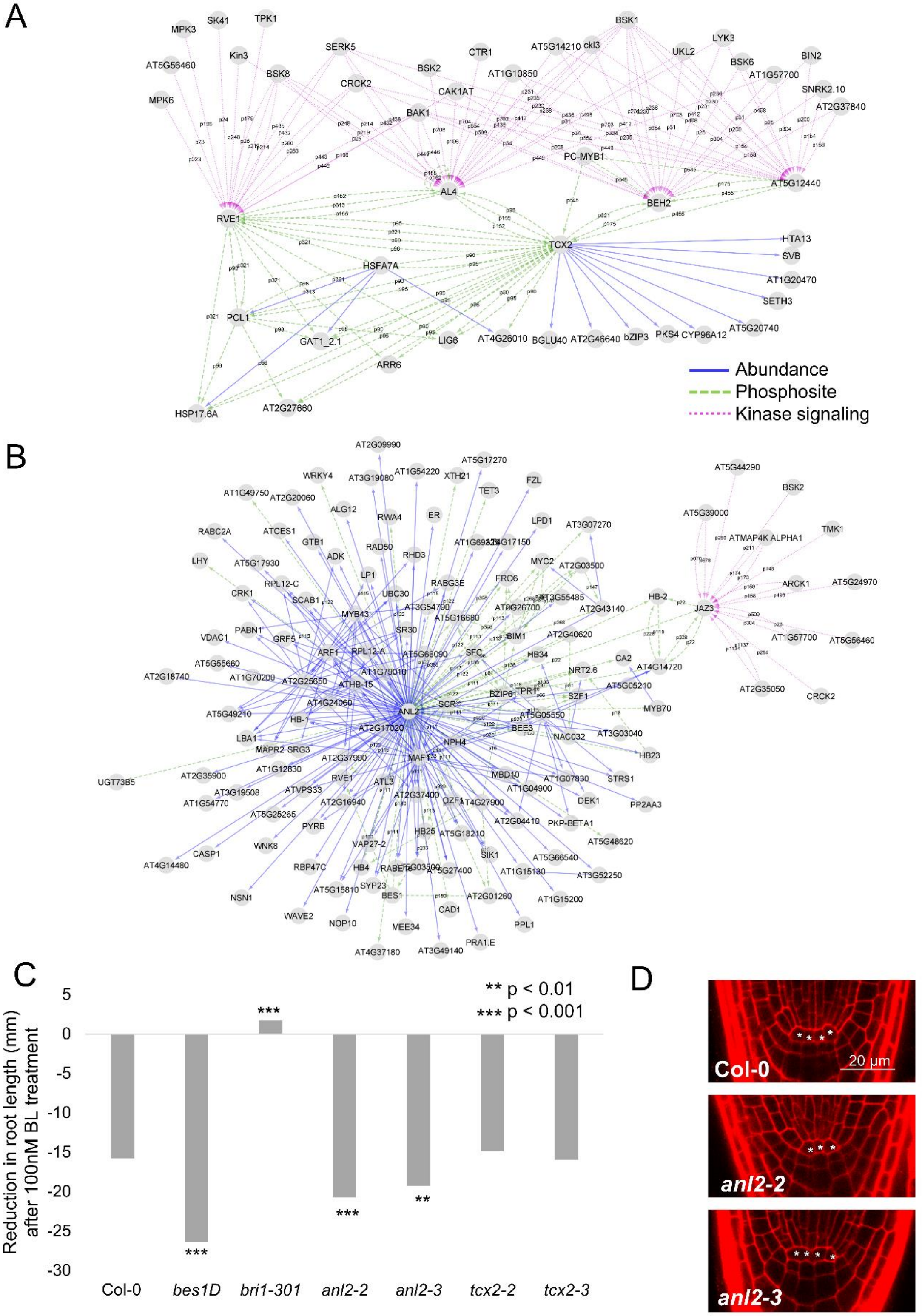

**Fig S4.** ANL2 is a BR-responsive TF. (A,B) First-neighbor subnetworks of TCX2 (A) and ANL2 (B). TF-abundance, blue solid; TF-phosphosite, green dashed; kinase signaling, purple dotted. For kinase-signaling, edge labels represent the phosphosite used to predict the regulation. (C) Response of *tcx2* and *anl2* mutant roots to 100 nM BL treatment. ** denotes $p < 0.01$, *** denotes $p < 0.001$, generalized mixed linear model. (D) Representative images of 5-day-old Col-0, *anl2-2* and *anl2-3* roots. * denotes QC cells.

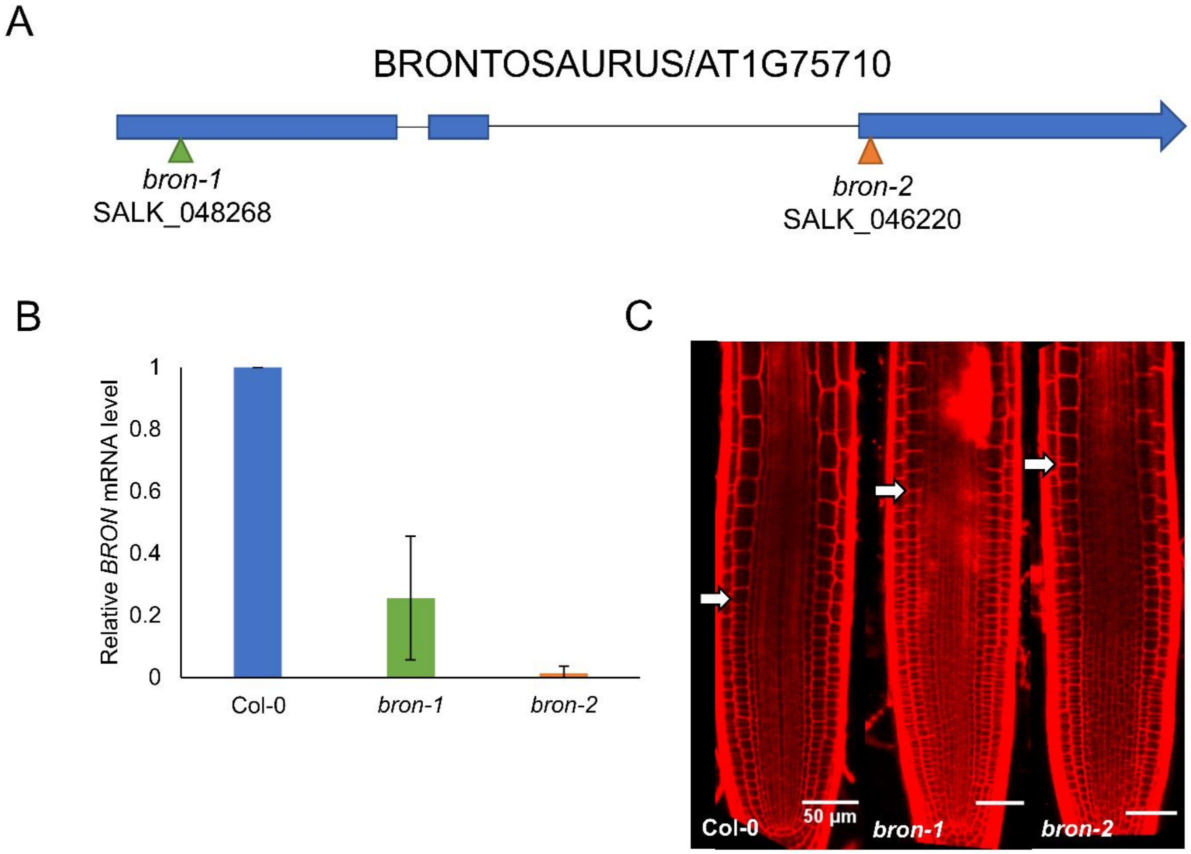

**Fig S5.** Two TDNA insertion alleles of BRONTOSAURUS. (A) Gene model of BRONTOSAURS showing locations of TDNA insertions. (B) RT-qPCR of BRONTOSAURS in Col-0 and the two TDNA insertion alleles. Bars represent SD. (C) Representative images of 5-day-old root meristems of Col-0, *bron-1* and *bron-2* mutants. Arrows mark the end of the meristem and the beginning of the transition zone.

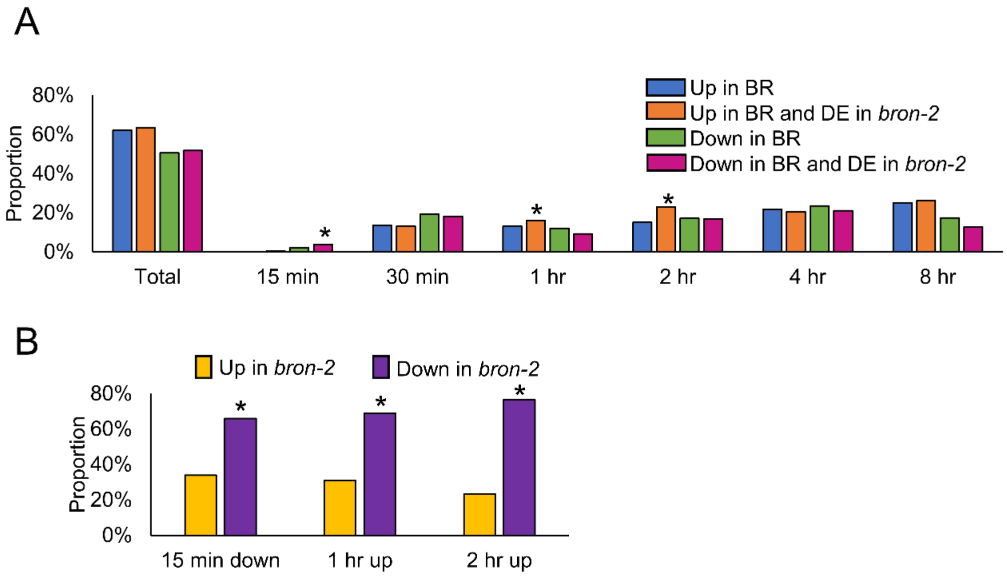

C

| GO term | Dn BR 15min & Up *bron-2* | Dn BR 15min & Dn *bron-2* | Up BR 1hr & Dn *bron-2* | Up BR 2hr & Dn *bron-2* |
|---|---|---|---|---|
| toxin metabolic process (GO:0009404) | ■ | | | |
| defense response (GO:0006952) | ■ | ■ | | |
| response to biotic stimulus (GO:0009607) | ■ | ■ | | |
| response to abiotic stimulus (GO:0009628) | ■ | ■ | ■ | ■ |
| response to fungus (GO:0009620) | ■ | ■ | | |
| response to stress (GO:0006950) | ■ | ■ | | |
| response to oxygen levels (GO:0070482) | | ■ | | |
| response to salicylic acid (GO:0009751) | | ■ | | |
| response to wounding (GO:0009611) | | ■ | | |
| immune response (GO:0006955) | | ■ | | |
| response to ethylene (GO:0009723) | | ■ | | |
| response to water deprivation (GO:0009414) | | ■ | | |
| response to hormone (GO:0009725) | | ■ | | |
| cell wall organization or biogenesis (GO:0071554) | | | ■ | |
| response to karrikin (GO:0080167) | | | ■ | ■ |
| response to temperature stimulus (GO:0009266) | | | ■ | ■ |
| flavonoid biosynthetic process (GO:0009813) | | | | ■ |
| response to heat (GO:0009408) | | | | ■ |
| response to UV (GO:0009411) | | | | ■ |
| response to stimulus (GO:0050896) | | | | ■ |

**Fig S6.** Overlap of BR responsive and BRON regulated genes. (A) Proportion of genes induced (blue) or repressed (green) by BR that are also differentially expressed in the *bron-2* mutant (orange and magenta, respectively). * denotes significant enrichment with $p<0.05$ using hypergeometric test. (B) Proportion of genes repressed by BR at 15 minutes or induced by BR at 1 or 2 hours that are up (yellow) or down (purple) in the *bron-2* mutant. * denotes $p<0.05$, Chi-squared test with likelihood ratio. (C) GO enrichment analysis of genes of interest. Green squares denote that term is enriched in that group of genes.

**Table S1. ChIP validation of predicted direct targets of BES1 from the integrated omics network**

| target | ChIP |
|---|---|
| AT1G13920 | NO |
| AT1G16610 | NO |
| AT1G20970 | NO |
| AT1G23380 | NO |
| AT1G23760 | NO |
| AT1G32360 | NO |
| AT1G32700 | YES |
| AT1G62300 | YES |
| AT1G71520 | NO |
| AT2G29480 | NO |
| AT2G30530 | YES |
| AT2G32730 | NO |
| AT2G33380 | NO |
| AT2G45660 | YES |
| AT2G47210 | NO |
| AT3G24020 | NO |
| AT3G26180 | NO |
| AT3G29075 | NO |
| AT3G44750 | NO |
| AT3G49620 | YES |
| AT3G51140 | NO |
| AT3G57570 | NO |
| AT5G10200 | NO |
| AT5G43830 | NO |
| AT1G16390 | NO |
| AT1G69030 | YES |
| AT2G15020 | NO |
| AT2G18160 | YES |
| AT2G29300 | NO |
| AT2G32560 | YES |
| AT2G34180 | NO |
| AT2G44430 | NO |
| AT3G14810 | NO |
| AT3G53620 | NO |
| AT4G03390 | NO |
| AT5G03520 | YES |
| AT5G47640 | YES |
| AT5G65613 | NO |
| AT1G01060 | NO |
| AT1G05300 | YES |
| AT1G09770 | NO |
| AT1G12740 | NO |

| | |
|---|---|
| AT1G13260 | YES |
| AT1G17840 | NO |
| AT1G30320 | YES |
| AT1G54120 | YES |
| AT1G58100 | YES |
| AT1G66200 | NO |
| AT1G69570 | YES |
| AT1G73390 | NO |
| AT1G76500 | YES |
| AT2G01260 | NO |
| AT2G23320 | YES |
| AT2G28930 | YES |
| AT2G37950 | NO |
| AT3G07570 | NO |
| AT3G25890 | NO |
| AT3G47500 | NO |
| AT3G47620 | YES |
| AT3G55980 | YES |
| AT4G00730 | YES |
| AT4G01810 | NO |
| AT4G18880 | YES |
| AT4G35750 | NO |
| AT4G36970 | NO |
| AT5G03470 | NO |
| AT5G14050 | NO |
| AT5G15130 | NO |
| AT5G17300 | NO |
| AT5G28300 | YES |
| AT5G38600 | NO |
| AT5G39660 | NO |
| AT5G41190 | NO |
| AT5G42920 | NO |
| AT5G49230 | NO |
| AT5G51440 | NO |
| AT1G49750 | NO |
| AT2G32540 | NO |
| AT3G02800 | NO |
| AT3G19450 | NO |
| AT3G48430 | NO |
| AT1G01540 | YES |
| AT1G04240 | YES |
| AT1G04430 | NO |
| AT1G06350 | NO |

| | |
|---|---|
| AT1G07370 | NO |
| AT1G10550 | YES |
| AT1G12460 | NO |
| AT1G15010 | NO |
| AT1G19968 | NO |
| AT1G29790 | NO |
| AT1G30690 | YES |
| AT1G34370 | NO |
| AT1G49380 | NO |
| AT1G55330 | YES |
| AT1G62300 | YES |
| AT1G62520 | YES |
| AT1G67750 | NO |
| AT1G72430 | YES |
| AT1G76160 | YES |
| AT2G19580 | YES |
| AT2G20760 | YES |
| AT2G26250 | NO |
| AT2G32690 | NO |
| AT2G33330 | NO |
| AT2G41820 | NO |
| AT2G41940 | YES |
| AT2G46710 | YES |
| AT2G47070 | YES |
| AT3G01970 | NO |
| AT3G02170 | YES |
| AT3G03850 | YES |
| AT3G05840 | YES |
| AT3G06770 | NO |
| AT3G07010 | YES |
| AT3G16910 | NO |
| AT3G23805 | NO |
| AT3G27650 | NO |
| AT3G28130 | NO |
| AT3G43720 | NO |
| AT3G46550 | YES |
| AT3G46940 | NO |
| AT3G57780 | NO |
| AT3G58790 | NO |
| AT4G00330 | NO |
| AT4G09160 | NO |
| AT4G14360 | NO |
| AT4G14440 | NO |

| | |
|---|---|
| AT4G14548 | YES |
| AT4G24275 | NO |
| AT4G29240 | NO |
| AT4G30800 | YES |
| AT4G34490 | NO |
| AT4G36240 | NO |
| AT5G01090 | YES |
| AT5G07870 | NO |
| AT5G16000 | NO |
| AT5G16590 | NO |
| AT5G18150 | YES |
| AT5G18690 | NO |
| AT5G19290 | NO |
| AT5G20130 | NO |
| AT5G23860 | YES |
| AT5G36260 | NO |
| AT5G41400 | YES |
| AT5G44670 | NO |
| AT5G46730 | NO |
| AT5G49170 | NO |
| AT5G51460 | NO |
| AT5G52280 | YES |
| AT5G54250 | NO |
| AT5G55960 | YES |
| AT5G56010 | NO |
| AT5G56860 | YES |
| AT5G62140 | NO |
| AT5G63180 | YES |
| AT5G64770 | NO |

**Table S2. BR phenotyping on mutants of interest**

| Line | BL0 Mean | BL100 Mean | Change in root length | p-val | Number of mock seedlings | Number of BR-treated seedlings |
|---|---|---|---|---|---|---|
| WT | 32.84 | 17.04 | -15.80 | | 73 | 73 |
| *bes1D* | 31.02 | 4.59 | -26.43 | 6.37E-06 | 28 | 28 |
| *bri1-301* | 26.36 | 28.06 | 1.69 | 5.71E-10 | 27 | 29 |
| *anl2-2* | 38.51 | 17.75 | -20.76 | 3.21E-03 | 40 | 34 |
| *anl2-3* | 31.60 | 12.35 | -19.25 | 1.33E-02 | 51 | 37 |
| *tcx2-2* | 36.77 | 21.87 | -14.90 | 5.71E-01 | 47 | 44 |
| tcx2-3 | 30.70 | 14.71 | -15.99 | 3.43E-01 | 46 | 42 |
| *bron-1* | 40.31 | 9.57 | -30.73 | 2.37E-19 | 48 | 39 |
| *bron-2* | 31.12 | 10.46 | -20.65 | 7.44E-04 | 43 | 54 |

**Table S3. TMT labeling strategy for quantitative proteomics**

| **BR1** | TMT label | Sample | Channel |
|---|---|---|---|
| | 126 | Ref | 1 |
| | 127N | Mock_15min_1 | 2 |
| | 127C | Mock_15min_2 | 3 |
| | 128N | Mock_15min_3 | 4 |
| | 128C | Mock_15min_4 | 5 |
| | 129N | BR_15min_1 | 6 |
| | 129C | BR_15min_2 | 7 |
| | 130N | BR_15min_3 | 8 |
| | 130C | BR_15min_4 | 9 |
| | 131 | Ref | 10 |
| **BR2** | TMT label | Sample | Channel |
| | 126 | Ref | 1 |
| | 127N | Mock_30min_1 | 2 |
| | 127C | Mock_30min_2 | 3 |
| | 128N | Mock_30min_3 | 4 |
| | 128C | Mock_30min_4 | 5 |
| | 129N | BR_30min_1 | 6 |
| | 129C | BR_30min_2 | 7 |
| | 130N | BR_30min_3 | 8 |
| | 130C | BR_30min_4 | 9 |
| | 131 | Ref | 10 |
| **BR3** | TMT label | Sample | Channel |
| | 126 | Ref | 1 |
| | 127N | Mock_1hr_1 | 2 |
| | 127C | Mock_1hr_2 | 3 |
| | 128N | Mock_1hr_3 | 4 |
| | 128C | Mock_1hr_4 | 5 |
| | 129N | BR_1hr_1 | 6 |
| | 129C | BR_1hr_2 | 7 |
| | 130N | BR_1hr_3 | 8 |
| | 130C | BR_1hr_4 | 9 |
| | 131 | Ref | 10 |
| **BR4** | TMT label | Sample | Channel |
| | 126 | Ref | 1 |
| | 127N | Mock_2hr_1 | 2 |

| | | | |
|---|---|---|---|
| | 127C | Mock_2hr_2 | 3 |
| | 128N | Mock_2hr_3 | 4 |
| | 128C | Mock_2hr_4 | 5 |
| | 129N | BR_2hr_1 | 6 |
| | 129C | BR_2hr_2 | 7 |
| | 130N | BR_2hr_3 | 8 |
| | 130C | BR_2hr_4 | 9 |
| | 131 | Ref | 10 |
| **BR5** | TMT label | Sample | Channel |
| | 126 | Ref | 1 |
| | 127N | Mock_4hr_1 | 2 |
| | 127C | Mock_4hr_2 | 3 |
| | 128N | Mock_4hr_3 | 4 |
| | 128C | Mock_4hr_4 | 5 |
| | 129N | BR_4hr_1 | 6 |
| | 129C | BR_4hr_2 | 7 |
| | 130N | BR_4hr_3 | 8 |
| | 130C | BR_4hr_4 | 9 |
| | 131 | Ref | 10 |
| **BR6** | TMT label | Sample | Channel |
| | 126 | Ref | 1 |
| | 127N | Mock_8hr_1 | 2 |
| | 127C | Mock_8hr_2 | 3 |
| | 128N | Mock_8hr_3 | 4 |
| | 128C | Mock_8hr_4 | 5 |
| | 129N | BR_8hr_1 | 6 |
| | 129C | BR_8hr_2 | 7 |
| | 130N | BR_8hr_3 | 8 |
| | 130C | BR_8hr_4 | 9 |
| | 131 | Ref | 10 |

**Table S4. Primers used in this study**

| Primer | Sequence |
|---|---|
| BES1 Fragment 1 Forward Primer | GCGGTACCATGACGTCTGACGGAGCAACGTCG |
| BES1 Fragment 1 Reverse Primer | TGGTGGAGTGACAGGAGCAGCGTTTGAGATTCTAAGTGG |
| BES1 Fragment 2 Forward Primer | CCTGTCACTCCACCAGTGGCAGCCCCAACTTCTAGAAACC |
| BES1 Fragment 2 Reverse Primer | CGCGGTACCACTATGAGCTTTACCATTTCCAA |
| BES1 Full Length Forward Primer | CGCGGTACCATGACGTCTGACGGAGCAACGTCG |
| BES1 Full Length Reverse Primer | CGCGGTACCACTATGAGCTTTACCATTTCCAA |
| BRON qPCR Forward Primer | AGGGTTTGTGCTTGTTCCCA |
| BRON qPCR Reverse Primer | GTTTCGACCCGAATCCTCCG |

**Supplemental Datasets**

**Dataset S1.** Transcripts differentially expressed in response to BR

**Dataset S2.** Protein groups differentially expressed in response to BR

**Dataset S3.** Phosphorylation sites differentially expressed in response to BR

**Dataset S4.** GO analysis on DE transcripts, protein groups, and phosphosites

**Dataset S5.** Inferred kinase-signaling and TF-centered networks

**Dataset S6.** Network Motif Analysis on TF-centered networks

**Dataset S7.** Transcripts differentially expressed in the *bron-2* mutant

**Dataset S8.** Overlap of transcripts DE in the *bron-2* mutant and the BR timecourse

**Dataset S9.** GO analysis on transcripts DE in the *bron-2* mutant and the BR timecourse.
